# SqueakOut: Autoencoder-based segmentation of mouse ultrasonic vocalizations

**DOI:** 10.1101/2024.04.19.590368

**Authors:** Gustavo M. Santana, Marcelo O. Dietrich

**Affiliations:** Laboratory of Physiology of Behavior, Interdepartmental Neuroscience Program, Program in Physics, Engineering and Biology, Yale University, USA; Graduate Program in Biochemistry, Federal University of Rio Grande do Sul, BRA; Laboratory of Physiology of Behavior, Department of Comparative Medicine, Department of Neuroscience, Yale University, USA

## Abstract

Mice emit ultrasonic vocalizations (USVs) that are important for social communication. Despite great advancements in tools to detect USVs from audio files in the recent years, highly accurate segmentation of USVs from spectrograms (i.e., removing noise) remains a significant challenge. Here, we present a new dataset of `12,954` annotated spectrograms explicitly labeled for mouse USV segmentation. Leveraging this dataset, we developed `SqueakOut`, a lightweight (`4.6M` parameters) fully convolutional autoencoder that achieves high accuracy in supervised segmentation of USVs from spectrograms, with a *Dice* score of `90.22`. `SqueakOut` combines a MobileNetV2 backbone with skip connections and transposed convolutions to precisely segment USVs. Using stochastic data augmentation techniques and a hybrid loss function, `SqueakOut` learns robust segmentation across varying recording conditions. We evaluate SqueakOut’s performance, demonstrating substantial improvements over existing methods like VocalMat (`63.82` *Dice* score). The accurate USV segmentations enabled by `SqueakOut` will facilitate novel methods for vocalization classification and more accurate analysis of mouse communication. To promote further research, we release the annotated `12,954` spectrogram USV segmentation dataset and the `SqueakOut` implementation publicly.

## 1 Introduction

Vocalizations are an important form of social communication among many animals, including mice [1]. Mice produce ultrasonic vocalizations (USVs) in various behavioral contexts such as maternal interactions [2, 3], social exploration [4], courtship [5], and distress situations like maternal separation [6, 7, 8, 9, 10, 11]. Although these high-frequency calls are inaudible to humans, they convey rich information about the animal’s internal state and behavioral experience [12, 13, 14]. For example, the emission of USVs by mouse pups when separated from their mother evokes maternal behavior and activates selective pathways in the brain of mothers [15, 16, 17, 18, 19]. Moreover, a detailed analysis of USVs is a powerful way to phenotype mouse models of neurodevelopmental disorders [20], genomic imprinting [16], pharmacological manipulations [21], as well as to perform comparative studies among different rodent species [22]. Thus, vocalizations provide a unique window into animals’ internal states and a rich framework for a better understanding of animal behavior and brain function.

The detailed analysis of USVs poses significant challenges due to the complexity of extracting precise spectrotemporal features. This general problem can be broadly divided into three components: detection, classification, and segmentation of USVs. Each component serves a specific purpose and presents unique challenges [23]. Detection involves identifying and distinguishing actual vocalization signals from periods of silence or noise in audio recordings (**Figure 1A)**. Accurate detection is crucial for the subsequent stages of analysis to be correct and meaningful (**Figure 1B)**. The detected USVs can be categorized into different classes, typically based on various acoustic features or specific characteristics identified on spectrograms of each vocalization [24]. Classification allows grouping similar sounds, aiding in studying the diversity and distribution of USVs across different contexts [1]. The third component is segmentation (**Figure 1B)**, which involves further breaking down each detected USV into discrete spectrotemporal units [25, 26]. Effective segmentation of USVs, akin to parsing words into distinct syllables, enables detailed analysis of the structure of each vocalization [27].

**Figure 1:**
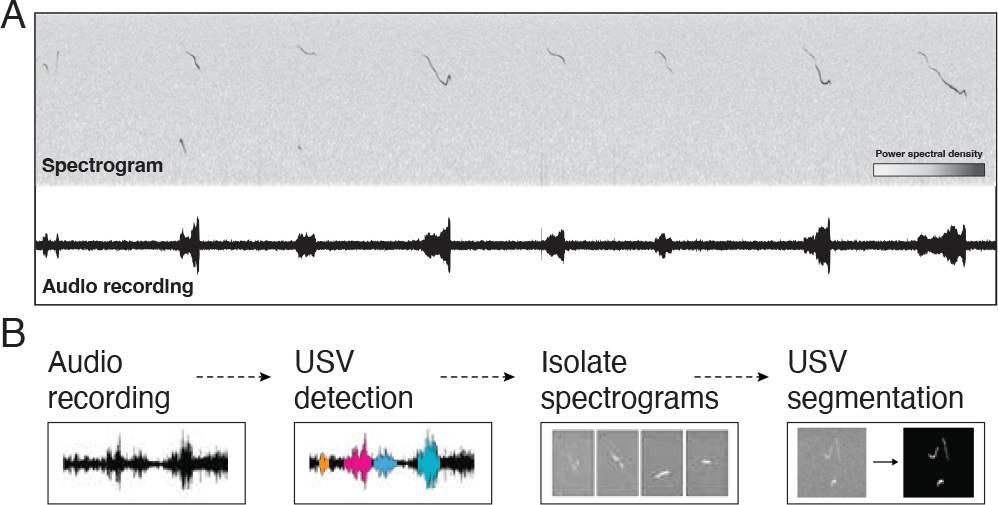
Overview of vocalization analysis pipeline. **(A)** The process begins with an audio recording, which is then converted into a spectrogram representation. **(B)** Individual vocalizations are detected, and isolated spectrograms are segmented and classified for downstream analysis.

Traditional methods for detecting and analyzing rodent USVs have relied heavily on manual annotation or semi-automated techniques based on predefined parameters and thresholds [28, 29]. These approaches are labor-intensive, time-consuming, and prone to human bias and error, especially when dealing with large datasets or complex vocal repertoires. Furthermore, they often fail to capture the nuanced spectrotemporal variations present in USVs, which may hold crucial information about the animal’s motivational state [13, 20]. Recent work has explored the application of computer vision techniques and machine learning models to USV audio recordings and spectrograms as a data-driven alternative [30, 26, 31, 26]. In particular, convolutional neural network (CNN) architectures have shown promising performance in detecting and classifying USVs [24, 30, 32]. Despite the significant progress made in the detection and classification of USVs, accurately segmenting USVs in spectrograms remains a challenge [30, 23, 31].

Here, we present a machine learning dataset explicitly designed for the segmentation of mouse USVs. Using this dataset, we also developed `SqueakOut`, a fully convolutional autoencoder for accurate USV segmentation. We evaluate `SqueakOut`’s performance, compare it to VocalMat, and demonstrate the utility of the segmented USVs for downstream analysis tasks. Lastly, we discuss insights gained from the model and potential applications to high-throughput USV phenotyping workflows.

## 2 Results

### 2.1 Creating a USV segmentation dataset

Creating a mouse USV segmentation dataset is a laborious task. One major problem is the intensive work required to produce accurate annotations by experts. Here, we generated an accurate segmentation dataset using both automated and manual approaches.

First, we took advantage of the publicly available dataset from VocalMat ([24]) and used it as a starting point (**Figure 2A**). The dataset consists of `12,954` spectrograms, including `2,083` spectrogram examples of noise and `10,871` spectrograms of mouse USVs. The dataset includes vocalizations from male and female mice of five different strains (C57Bl6/J, NZO/HlLtJ, 129S1/SvImJ, NOD/ShiLtJ, and PWK/PhJ), ranging from postnatal day 5 to postnatal day 15.

**Figure 2:**
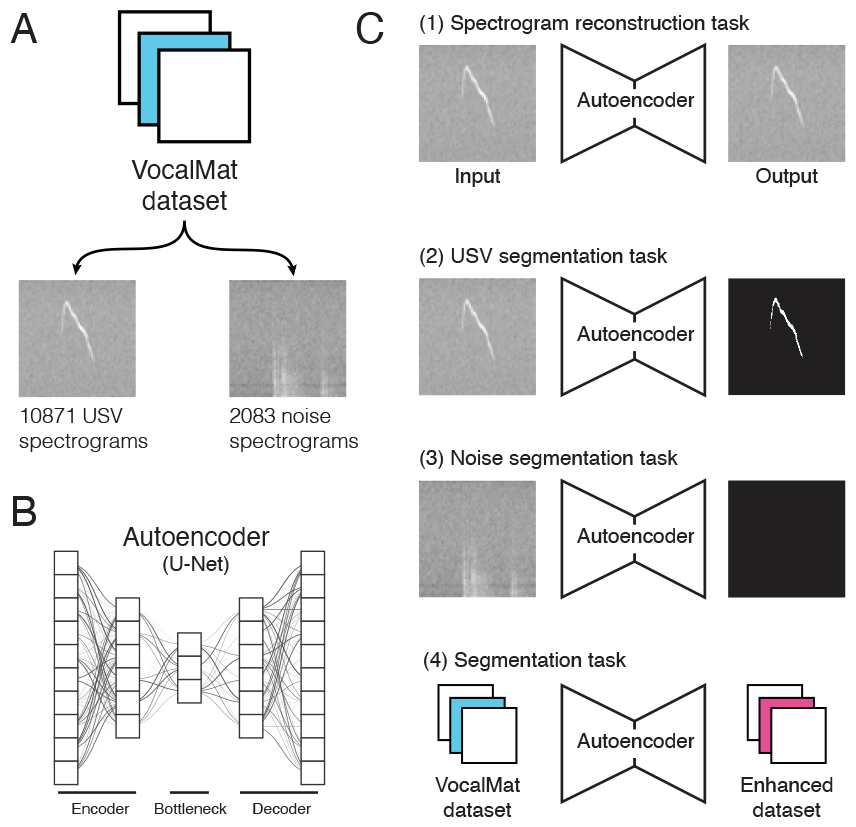
Automated methods for creating the USV segmentation dataset. We use the VocalMat dataset [24] which contains 10,871 USV spectrograms and 2,083 noise spectrograms **(A)**, and the U-Net autoencoder architecture [33] **(B)**. The autoencoder is trained on a spectrogram reconstruction task **(C1)**. The pre-trained autoencoder is then trained on a segmentation task using USV spectrograms **(C2)** and on noise spectrograms **(C3)**. The trained U-Net model is used to enhance the original VocalMat dataset by generating segmentation masks that are mostly devoid of noise components **(C4)**.

#### 2.1.1 Automated approaches for creating a USV segmentation dataset

VocalMat uses computer vision and machine learning to detect, segment, and classify vocals. While VocalMat’s USV detection rate achieves state-of-the-art results, its segmentation performance is suboptimal when noise is present in spectrograms (see **Figure 3C**), making the segmentation masks unsuitable for directly training a segmentation neural network. To enhance the VocalMat dataset, we used its segmentation masks as a starting point and trained an autoencoder to learn the segmentation task. We trained the autoencoder separately on spectrograms of USVs and noise to enable it to learn representations of vocalizations and noise. This approach allowed us to use the autoencoder to denoise the original segmentation masks. The following automated processing steps result in a dataset with segmentation masks of vocalizations with fewer noise segments.

**Figure 3:**
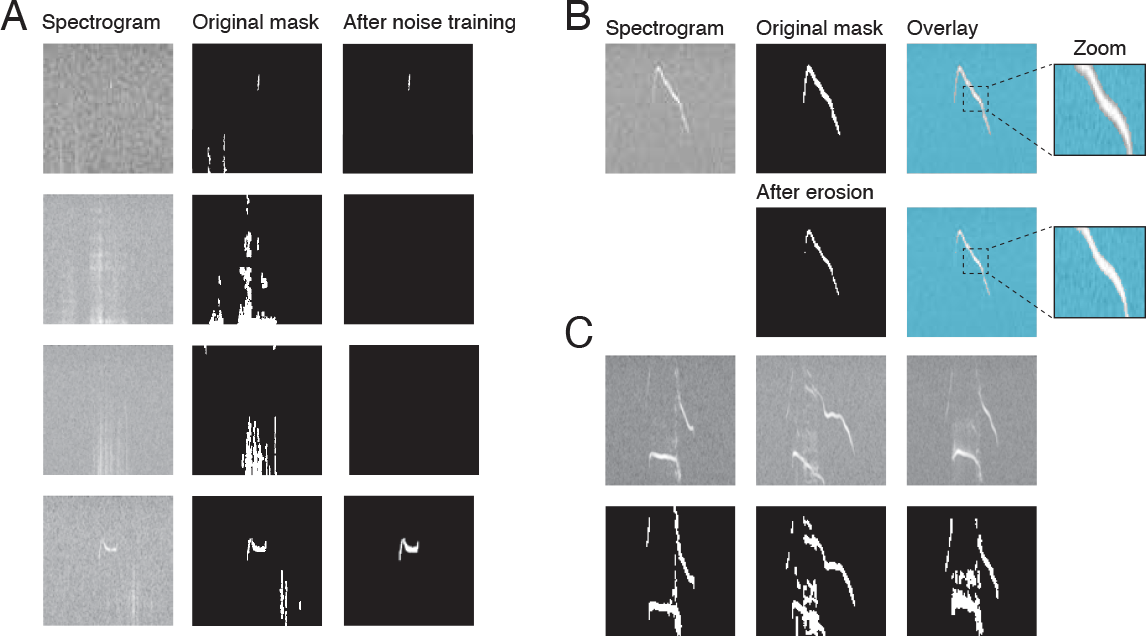
Dataset fine-tuning processing. **(A)** Comparison of an original VocalMat noisy mask and the trained autoencoder mask. **(B)** Illustration of the *border thinning* process through erosion operation applied to a segmentation mask. **(C)** Three examples illustrating noise overlapping with USVs in spectrograms and the segmentation produced by VocalMat, highlighting the challenges in accurate vocalization segmentation.

##### Unsupervised spectrogram reconstruction task

We begin by using U-Net, an autoencoder broadly used for biomedical image segmentation [33] (**Figure 2B**), and train the network on the VocalMat dataset for the unsupervised task of spectrogram reconstruction (**Figure 2C-1)**. In this step, the autoencoder receives the spectrograms as input and is trained to reconstruct the input spectrogram as its output. The autoencoder has a bottleneck architecture to ensure that it learns a meaningful representation of the data instead of simply copying its input as the output. The autoencoder can reconstruct spectrograms with high accuracy (`91.67`% *±* `2.08`%; mean *±* SEM).

##### USV segmentation task

Next, using the pre-trained autoencoder from the previous step, we trained the autoencoder for the task of spectrogram segmentation (**Figure 2C-2**). The autoencoder receives spectrograms of USVs as inputs and outputs binary segmentation masks, i.e., images containing `0` and `1`, where `1` indicate USV segments. Initially, we use the segmentation produced by VocalMat as the ground truth annotations for this training step.

##### Noise segmentation task

Similarly to the previous step, we trained the autoencoder for the task of spectrogram segmentation but only using the spectrograms of noise (**Figure 2C-3**). This allows the autoencoder to learn and distinguish between representations of noise and actual vocalizations in the spectrograms.

##### Enhancing the dataset

Following the pre-training steps, we now use the autoencoder to perform inference over the original VocalMat dataset for the task of spectrogram segmentation (**Figure 2C-4**). Since the autoencoder has learned the task of spectrogram segmentation and representations of noise, it can produce segmentation masks similar to those generated by VocalMat but with fewer false positives (spectrograms containing only noise: VocalMat `2.85`% *±* `1.27`%; Autoencoder `78.91`% *±* `4.86`%; values are the *Dice* score mean *±* SEM) (**Figure 3A**).

#### 2.1.2 Manual fine-tuning of the USV segmentation dataset

The generated dataset following the automated steps is significantly better than the original dataset (**Figure 3A**) but still requires fine-tuning. VocalMat’s algorithms produce segmentation masks that exceed the size of the actual vocalizations, capturing both vocal-related and surrounding pixels. (**Figure 3B**). We address this *border effect* by using the morphological image processing technique known as erosion. This thinning process effectively reduces the size of the segmentation masks, mitigating the *border effect* (**Figure 3B**). We apply erosion to all vocalization segments at least four times as large as the erosion kernel (`4`*×*`2` pixels) to prevent the thinning out of already small vocalizations.

Following the *border* thinning process, we repeat automated steps `(2)` through `(4)`. This sequence of automated and manual refinements is repeated until erosion can no longer be applied to any vocalization segments.

The final step in creating the dataset involves manual refinement. Despite having over two thousand noise examples from the VocalMat dataset, certain types of noise are exclusively present in spectrograms containing vocalizations. Consequently, these noise segments are incorrectly detected as vocalizations (**Figure 3C**). An expert annotator manually inspected all segmentation masks using the image annotation and segmentation tool `RectLabel`^1^, and corrected the masks for a small subset (approximately 15%) of the data.

### 2.2 SqueakOut: Autoencoder for mouse USV segmentation

#### 2.2.1 Network architecture

`SqueakOut` is a fully convolutional autoencoder that generates segmentation masks of vocalizations from spectrograms. The architecture is depicted in **Figure 4**. `SqueakOut` uses a modified MobileNetV2 [34] as its backbone for its small memory footprint and efficient processing using inverted residuals, depth-wise convolutions, and lack of explicit non-linearities in the narrow layers. Specifically, we removed the *average pooling* layer and added a *dropout* layer before the final bottleneck layer with a 20% rate.

**Figure 4:**
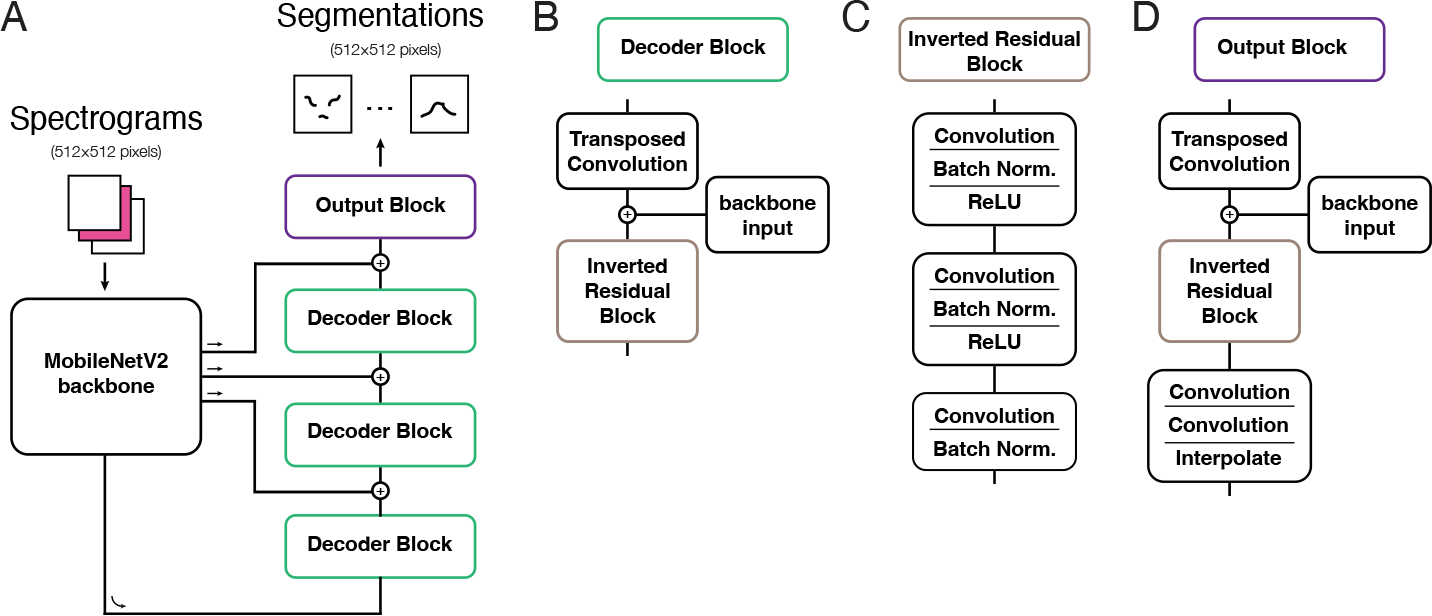
SqueakOut network architecture combines a MobileNetV2 backbone with an encoderdecoder structure. **(A)** SqueakOut takes input spectrograms and outputs segmentations masks (both `512`*×*`512` pixels). Spectrograms are processed through the backbone encoder, and then through a series of decoder blocks (shown in **B** and **C**) with skip connections and transposed convolutions and an output block (shown in **D**) to produce segmentation masks.

The decoder path for `SqueakOut` uses skip connections from backbone layers and transposed convolutions to reconstruct segmentation masks. Skip connections have been demonstrated to enhance segmentation accuracy and capture fine-grained details [35, 36, 37]. Moreover, skip connections improve gradient propagation and convergence during training [37, 38]. In `SqueakOut`, we concatenate the input from backbone layers with the input from the previous layer in the decoder path, ensuring the propagation of intact information from early layers in the network. Lastly, we employ a series of convolutional layers in the output block and use interpolation to upsample the output, restoring it to the same spatial dimensions as the original input spectrogram.

#### 2.2.2 Training

`SqueakOut` is implemented in PyTorch [39] using the PyTorchLightning framework [40] and was trained on the enhanced VocalMat dataset. A subset of the dataset containing `849` USVs was used as the *test* set. The remaining dataset was randomly split into *training* (90%) and *validation* (10%) sets. `SqueakOut` was trained using Adam [41] with a learning rate of 1*e*^−4^. The learning rate was reduced by a factor of 0.1 if performance on the *validation* set did not improve for five consecutive iterations. Training was halted if the performance did not improve for fifteen consecutive iterations to avoid overfitting. A batch size of eight samples was used. The loss function was a weighted sum of the Focal loss (FL) [42]—which emphasizes hard data points and prevents easy negatives from dominating the loss during training—, and the Dice loss (DL) [43]—a measure of the similarity between the network output and the ground-truth segmentation. We chose this hybrid loss function because of the extreme imbalance in class labels for the segmentation task. The majority of pixels in a spectrogram represent background, with USVs accounting for, on average, less than `5`% of pixels. Briefly,

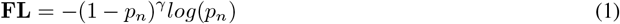

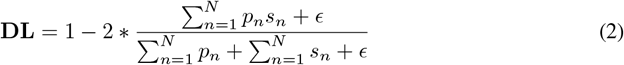

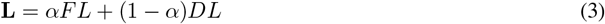

where *p*_*n*_ is the predicted segmentation probability map, and *s*_*n*_ is the ground-truth segmentation map for a spectrogram. We include a small term *ϵ* in the *Dice* loss to prevent division by `0`. In `SqueakOut`, we use *γ* = 2 and *α* = 0.3.

#### 2.2.3 Data augmentations

To further enrich our dataset, we utilized data augmentation techniques. The segmentation task is unique in that any augmentations made on the spectrogram can be identically applied to its matching segmentation mask. Importantly, this would not hold for a classification task. For example, if a spectrogram is randomly warped such that the morphological characteristics of a USV changes, then the corresponding classification of that USV would also likely change in unpredictable ways.

To make `SqueakOut` robust to changes in spectrogram quality and better generalize, we use the data augmentations depicted in **Figure 5**. Briefly, augmentations were applied during training on a batch-by-batch basis with the following conditions:

**Figure 5:**
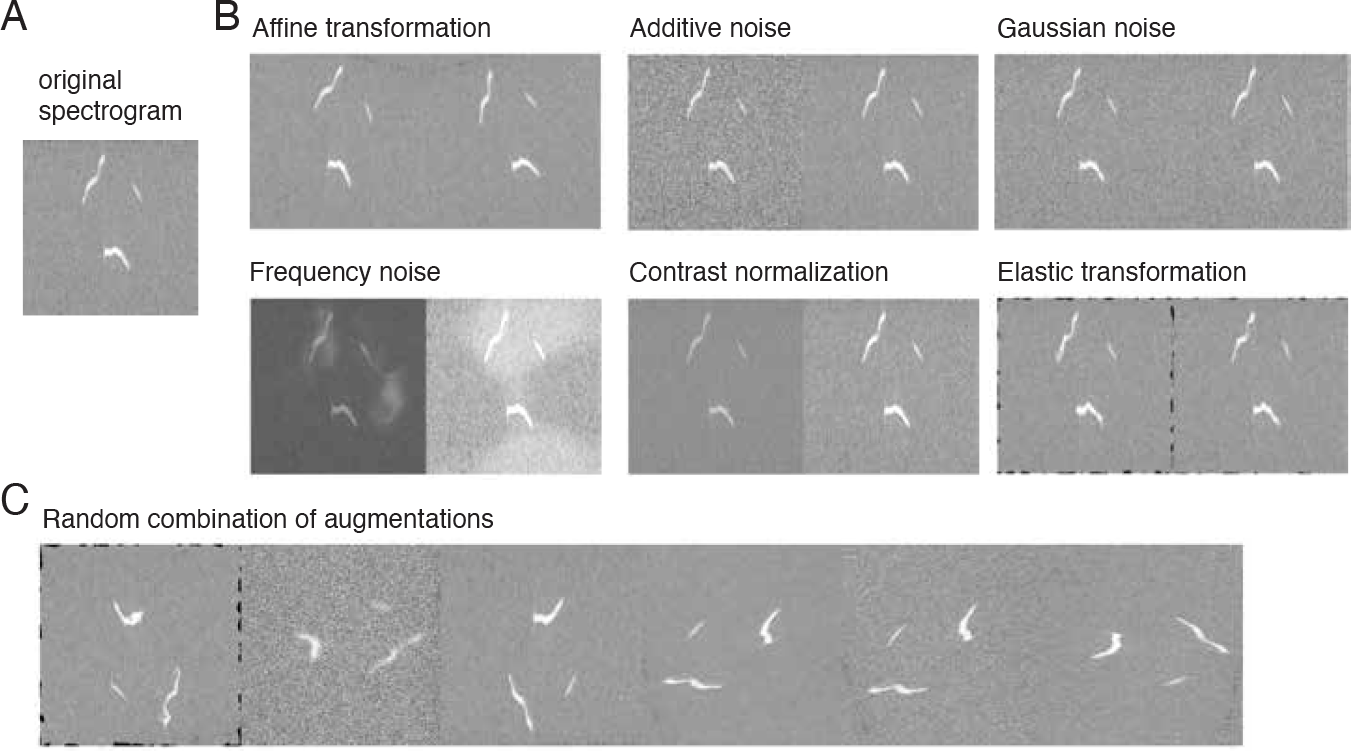
Dataset augmentation techniques applied to spectrograms for training robust segmentation models. **(A)** An example USV spectrogram. **(B)** Illustration of individual data augmentation techniques applied to the example spectrogram. **(C)** Illustration of augmentation techniques when jointly applied to the example spectrogram.

a. 75% chance of applying a random affine transformation
b. 25% chance of applying contrast normalization
c. 25% chance of applying a Gaussian blur
d. 33% chance of applying one of the following:

- Additive noise
- Gaussian noise
- Frequency noise
- Elastic transformations

Augmentations for each of the four conditions (*a*-*d*) have an independent probability of being applied to a given batch of spectrograms. This means that for each training batch, any combination of the four augmentations can occur, ranging from no augmentations to all four being applied simultaneously. The stochastic nature of this augmentation strategy helps to increase the diversity of the training data and improve `SqueakOut`’s robustness to variations in the input spectrograms. For example, noise and contrast augmentations especially improve segmentation in spectrograms with low signal-to-noise ratios. Affine and elastic transformations are intended to make `SqueakOut` robust to USV morphologies not present in our dataset and improve USV contour segmentation quality.

### 2.3 USV segmentation performance

`SqueakOut` is a lightweight autoencoder model at only `18MB` (`4.6M` parameters) that is fast and achieves high accuracy. Inference on a batch of `64 512`*×*`512 pixels` spectrograms on a gaming GPU takes less than `0.035` seconds, and about `8` seconds on a CPU.

Vocalizations on spectrograms are generally small, leading to an imbalance in class labels (`0` or `1`) for the segmentation task. We therefore created a `null model`, which treats every spectrogram as if it contains no USVs, i.e., it always generates blank segmentation masks (all values are `0`). The `null model` achieves `99.09`% accuracy, showing that most pixels in a spectrogram are indeed background. Similarly, all models achieve relatively high pixel-wise accuracy (**Table 1**). In contrast, the null model achieves a `4.95` *Dice* score. The *Dice* score measures the overlap between two segmentation masks by weighing their intersection against their union (**Equation 2**), where a score of `100` means perfect overlap. It effectively balances false positive and false negative rates, and, therefore, we use the *Dice* coefficient as our primary performance score metric.

**Table 1:** USV segmentation performance across models. Values are mean *±* SEM.

| Model | Performance |  |
| --- | --- | --- |
|  | Accuracy (%) | Dice score |
| Null Model | 99.10 ± 0.02 | 4.95 ± 0.74 |
| VocalMat [24] | 98.83 ± 0.06 | 63.82 ± 0.71 |
| U-Net [33] | 98.91 ± 0.04 | 72.31 ± 0.83 |
| SqueakOut (no augmentations) | 99.03 ± 0.02 | 82.98 ± 0.65 |
| SqueakOut | 99.84 ± 0.01 | 90.22 ± 0.41 |

We first compared `SqueakOut`’s performance with VocalMat and applied the same metrics to the output of both tools (**Table 1**). VocalMat is accurate in segmenting any high-intensity segments in spectrograms, including noise (**Figure 6**), resulting in a significantly lower *Dice* score. In noise-free recordings, we expect that VocalMat’s performance would be qualitatively similar to `SqueakOut`. We also compared `SqueakOut` with another autoencoder architecture for image segmentation, U-Net. We trained the U-Net model and `SqueakOut` using the same dataset, without any data augmentations. Nevertheless, the U-Net model performs worse than `SqueakOut` (no augmentations) (**Table 1**), showing that `SqueakOut`’s high *Dice* score is not solely due to our refined dataset, but also to its architecture. Importantly, the data augmentation techniques we employed improved `SqueakOut`’s *Dice* score by `8.72`% (last two rows in **Table 1**).

**Figure 6:**
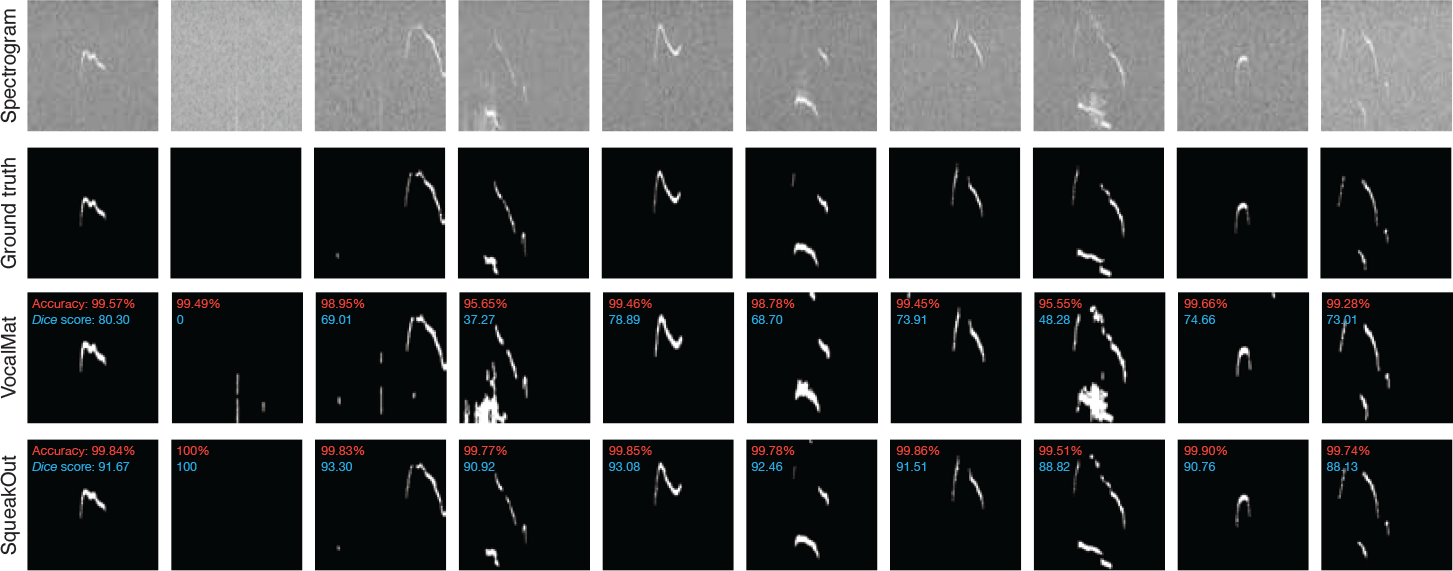
Comparison of USV spectrogram segmentation performance between the ground truth annotations, the VocalMat dataset, and the proposed `SqueakOut` method. The spectrograms illustrate `SqueakOut`’s ability to accurately segment USVs compared to the ground truth and the baseline VocalMat dataset. Inset values in red represent the pixel-wise accuracy, while those in blue indicate the *Dice* score.

## 3 Discussion

Here, we first present a new mouse USV segmentation dataset that is made publicly available and can be used by any group for machine learning applications. We leveraged this unique dataset to develop `SqueakOut`, a fully convolutional autoencoder for supervised segmentation of USV spectrograms. Our results demonstrate `SqueakOut`’s ability to segment USVs with very high accuracy. Our segmentation dataset joins the VocalMat dataset to provide a single high-quality annotated dataset for mouse USV detection and segmentation.

Autoencoders have been used across a broad spectrum of research in various species to analyze acoustic communication [44, 45, 30, 46, 47]. The popularity of autoencoders is partly due to their inherent ability to learn latent data structures in an unsupervised fashion, i.e., without requiring annotated datasets. This can be advantageous when the features of the data relevant for analysis are unknown or for unbiased data analysis. However, the extracted latent features are often hard to interpret. On the other hand, supervised methods that use pre-defined features are interpretable but can lead to biased analysis depending on the features chosen by the experimenter.

Unsupervised methods that attempt to learn latent features of vocals using their spectrograms often suffer from variability in the quality of recordings. Any variability in recording conditions will result in drastically different background noise levels and signal-to-noise ratios in spectrograms. This variability will affect what the network learns and can make latent features even harder to interpret. Unsupervised methods have to be carefully tuned to the specifics of a dataset, and an extensive list of methods have been developed to deal with varying quality in spectrograms [47, 23]. `SqueakOut` can produce accurate USV segmentations, effectively removing any variability due to recording conditions. The resulting segmentation masks can be used for downstream analysis using unsupervised methods such as Variational Autoencoders [45, 30] and dimensionality reduction techniques such as UMAP [48] or diffusion maps [49] to exploit the combined advantages of unsupervised and supervised methods.

Vocalizations in mice are powerful indicators of their emotional and behavioral states, and classification of these vocalizations is important for linking behavior with brain function across different contexts [1]. Historically popular methods have used hand-crafted features such as the duration, frequency modulation, amplitude, and other characteristics of each USV. These methods heavily relied on the quality of the segmentation and were therefore not incredibly precise nor high-throughput. Recently, CNNs have become the standard models for supervised image classification and are widely used for USV call type classification but lack spatiotemporal measurements (e.g., USV duration or average frequency). However, with accurate segmentations such as those produced by `SqueakOut`, the use of hand-crafted features and traditional machine learning methods such as random forests can reemerge as efficient and powerful alternatives for studying USV diversity across behaviors using interpretable features.

In conclusion, this work presents a new publicly available dataset for mouse USV segmentation, which we believe will be a valuable resource for the research community. We demonstrate the utility of this dataset by developing `SqueakOut`, a fully convolutional autoencoder that achieves high accuracy in supervised USV segmentation. The combination of our segmentation dataset with the existing VocalMat dataset provides a comprehensive, high-quality annotated resource for USV detection and segmentation. By providing accurate segmentation tools, we aim to enable more precise and powerful methods for USV classification and analysis, facilitating novel approaches for studying mouse communication and neurobiology.

## 4 Materials and methods

### Mouse USV dataset

The annotated dataset for mouse USV segmentation is openly available on Open Science Framework [50] at https://osf.io/f9sbt/. We welcome contributions from the community. Anyone may submit corrections or newly annotated audio recordings to be included in the dataset. A similar approach to this work can be used to generate segmentation labels.

### SqueakOut architecture

SqueakOut was implemented in PyTorch `v1.7.0` [39]. The network implementation is available at https://github.com/gumadeiras/squeakout. Here we provide a brief overview of the PyTorch functions utilized to implement each layer: convolution (`Conv2d`), batch normalization (`BatchNorm2d`), ReLU6 (`ReLU6`), dropout (`Dropout`), transposed convolution (`ConvTranspose2d`), and upsampling (`Interpolate`).

### Pre-trained SqueakOut network

The SqueakOut network implementation and pre-trained weights are available at https://github.com/gumadeiras/squeakout. We provide scripts to train SqueakOut on a new dataset or perform inference on your data. We developed and have tested SqueakOut using Python `3.7.10`, NumPy `v1.21.5` [51], Scikit-Learn `v1.0.2` [52], PyTorch `v1.7.0` [39], Torchvision `0.8.0`, PyTorchLightning `v1.4.0` [40], and the image augmentation library `imgaug v0.4.0`.

### Segmentation metrics

We used two metrics to quantify segmentation performance: pixel-wise accuracy, and the *Dice* score. Pixel-wise accuracy was computed as follows:

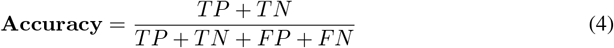

where *TP* are the true positives (correctly predicting a pixel belongs to a vocalization), *TN* the true negatives (correctly predicting a pixel belongs to the background), *FP* the false positives (incorrectly predicting a pixel belongs to a vocalization), and *FN* the false negatives (incorrectly predicting a pixel belongs to the background). The *Dice* score was computed as described in **Equation 2**.

## Acknowledgments and Disclosure of Funding

We thank Antonio Fonseca for insightful discussions on the early development of the segmentation dataset. We thank Sarah Mohr for suggesting the name `SqueakOut`. M.O.D. was supported by the National Institute of Mental Health of the National Institutes of Health (R01MH125008 and R01MH130825), by the Smith Family Foundation, and by discretionary funds from the Yale School of Medicine. G.S. was supported in part by the Coordination for the Improvement of Higher Education Personnel – Brazil (CAPES) – Finance Code 001.

## Footnotes

1 https://rectlabel.com

